# Immunogenicity and efficacy of a subcutaneously administered, adjuvanted vaccine containing modified S1 spike protein of SARS-CoV-2 variant C.1.2

**DOI:** 10.1101/2023.08.31.555805

**Authors:** Michael Bowe, Emily Wright, Peter Pushko, Brian Backstedt, Casey Gardner, Micah Short-Freeman, Swagata Kar, David Craig Wright

## Abstract

During the COVID-19 pandemic, vaccines have produced protective immunity sufficient enough to cause a decrease in hospitalizations and deaths; however, the pandemic continues due to mutational events, predominantly occurring in the S1 sequence of the spike protein of SARS-CoV-2. We have developed a baculovirus-expressed, modified S1 SARS-CoV-2 protein based on the C.1.2 variant, which was first identified in South Africa.^1^ This was encapsulated in a vitamin E containing, nonphospholipid liposome, which was then used to subcutaneously immunize Syrian hamsters. This vaccine, when administered at day 1 generates IgG responses that react to the modified C.1.2 S1 protein; full-length spike proteins from Wuhan-Hu-1, Delta, Omicron BA.1; and the Omicron recombinant variant XBB.1.5 in 100% of the animals. The second dose administered subcutaneously on day 28 demonstrated anamnestic response in the quantitative IgG assay to the Wuhan-Hu-1 spike Receptor Binding Domain (RBD). In addition, antibody IgA and IgM responses in sera were demonstrated. Serum IgG antibody responses to the spike proteins of the modified C.1.2 S1 and full-length spike proteins Wuhan-Hu-1, Delta, Omicron BA.1, and Omicron recombinant XBB.1.5 variants are elevated for over 120 days. Challenge of vaccinated and unvaccinated hamsters at day 126 of the study with an Omicron BA.1 resulted in a difference in weight change and viral load based on the qRT-PCR assay seven days after challenge.

## 1. Introduction

The majority of FDA-approved vaccines are administered intramuscularly rather than subcutaneously. An exception to intramuscular injection of vaccines is the inactivated polio vaccine (Sanofi) which is also FDA-approved for subcutaneous administration. This approval raises the interesting possibility that one could develop other adjuvanted vaccines that could be administered subcutaneously. The utility of this approach is that no muscle damage would occur from the subcutaneously injected vaccine and, presumptively, one would have no pain and reduced local swelling of the arm. In addition, there should be no circulating spike protein in the vascular space as is seen in SARS-CoV-2 infection and mRNA vaccination.^2^ The occurrence of the SARS-CoV-2 pandemic provided us the opportunity to evaluate this hypothesis with an adjuvanted, modified S1 sequence of the spike protein of the C.1.2 variant of SARS-CoV-2. We chose to evaluate spike proteins from two isolates discovered in Africa in May 2021, the C.1.2 isolate^3^ and the Omicron BA.1 isolate. This paper discusses the work with our modified S1 sequence of the spike protein of C.1.2. We have designed and produced negatively charged, fusogenic, vitamin E containing nonphospholipid liposomes^4, 5, 6, 7^ containing the modified S1 spike protein of isolate C.1.2. This variant was chosen for our antigen as it contained many of the mutations that had caused worldwide infections and death. Table 1 below shows the mutations present in the spike protein from candidate isolates of Delta, BA.1, and C.1.2.

**Table 1.** Mutations present in the spike protein from candidate isolates of Delta, BA.1, and C.1.2 compared to S1 construct.

| Delta | BA.1 | BA.1 (cont) | C.1.2 | S1 Construct |
| --- | --- | --- | --- | --- |
| T19R | A67V | P681H | P9L | P9L |
| E156G | del69/70 | N764K | C136F | P25L |
| del157/158 | T95I | D796Y | Del144Y | C136F |
| L452R | G142D |  | R190S | Del144Y |
| T478K | del143/145 |  | D215G | W152K |
| D614G | N211I |  | Del243-244AL | R190S |
| P681R | del212/212 |  | Y449H | D215G |
| D950N | G339D |  | E484K | Del243-244AL |
|  | S371L |  | N501Y | F275Y |
|  | S373P |  | D614G | L276I |
|  | S375F |  | H655Y | R346K |
|  | S477N |  | N679K | N440K |
|  | T478K |  | T716I | Y449H |
|  | E484A |  | T859N | E484K |
|  | Q493R |  |  | N501Y |
|  | G496S |  |  | D614G |
|  | Q498R |  |  | H655Y |
|  | N501Y |  |  | N679K |
|  | Y505H |  |  | P681H |
|  | T547K |  |  |  |
|  | D614G |  |  |  |
|  | H655Y |  |  |  |
|  | N679K |  |  |  |

In this hamster study, we demonstrated the ability of our adjuvanted, modified S1 vaccine construct in our vitamin E containing nonphospholipid liposome to generate IgM, IgA, and IgG antibodies to multiple variants of SARS-CoV-2 and cross-protection from an Omicron BA.1 isolate in a hamster model, when challenged 126 days after primary vaccination.

## 2. Materials and Methods

### 2.1 Design of the C.1.2 S1 construct

In April 2020, Lyons-Weiler identified all of the homologous peptide sequences between SARS-CoV-2 and human proteins and found five sequences containing peptides with homology to seven human proteins.^8^ Based on this information, we decided to delete amino acids 683-1,270 of the full-length spike protein from our modified S1 construct. In deleting amino acids after position 683 of the spike protein, five sequences containing peptides with homology to seven human proteins have been removed. These seven proteins (and their sites in the human body) are keratin associated protein 4-7 (KRTAP4-7) (skin), Metallothionein 1 E (MT1E) (liver and other tissues), human coiled-containing domain-containing protein 175 isoform X8 (brain, pituitary gland, and testis), follistatin-related protein 1 isoform X1 (placenta), tetratricopeptide repeat protein 28 isoform X8 (ubiquitous in human tissue), ALDH1L1 protein (ubiquitous in human tissue), and attractin-like protein 1 (brain). See Table 2 for additional details.

**Table 2.**
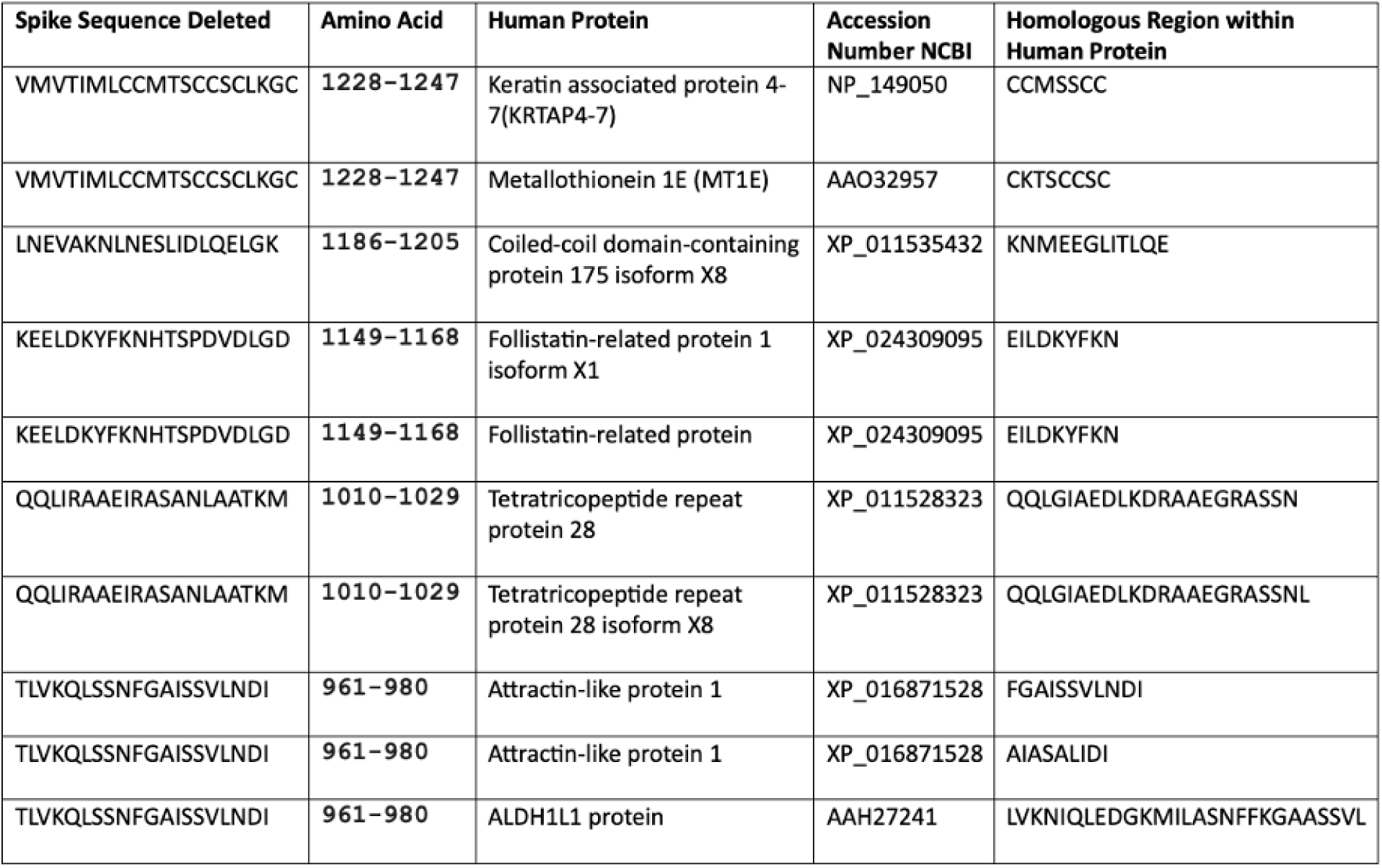
SARS-CoV-2 spike sequences with homology to human proteins.

### 2.2 Preparation of modified C.1.2 S1 protein and expression of genes of interest (GOI), modified S1 sequence of C.1.2 (Figure 2) in baculovirus expression vector system

#### 1. Synthesis and cloning of GOI into transfer vector

The GOI sequence (Figure 2) was based on the modified S1 sequence of the spike protein of C.1.2 (Figure 1) provided by D4 Labs, LLC. The gene was designed by optimization of nucleotide codon bias for high-level expression in insect cells. The gene was biochemically synthesized de novo and cloned into pFastBac1-baculovirus transfer vector. The GOI sequence and adjacent vector sequences were confirmed by DNA sequencing.

**Figure 1.**
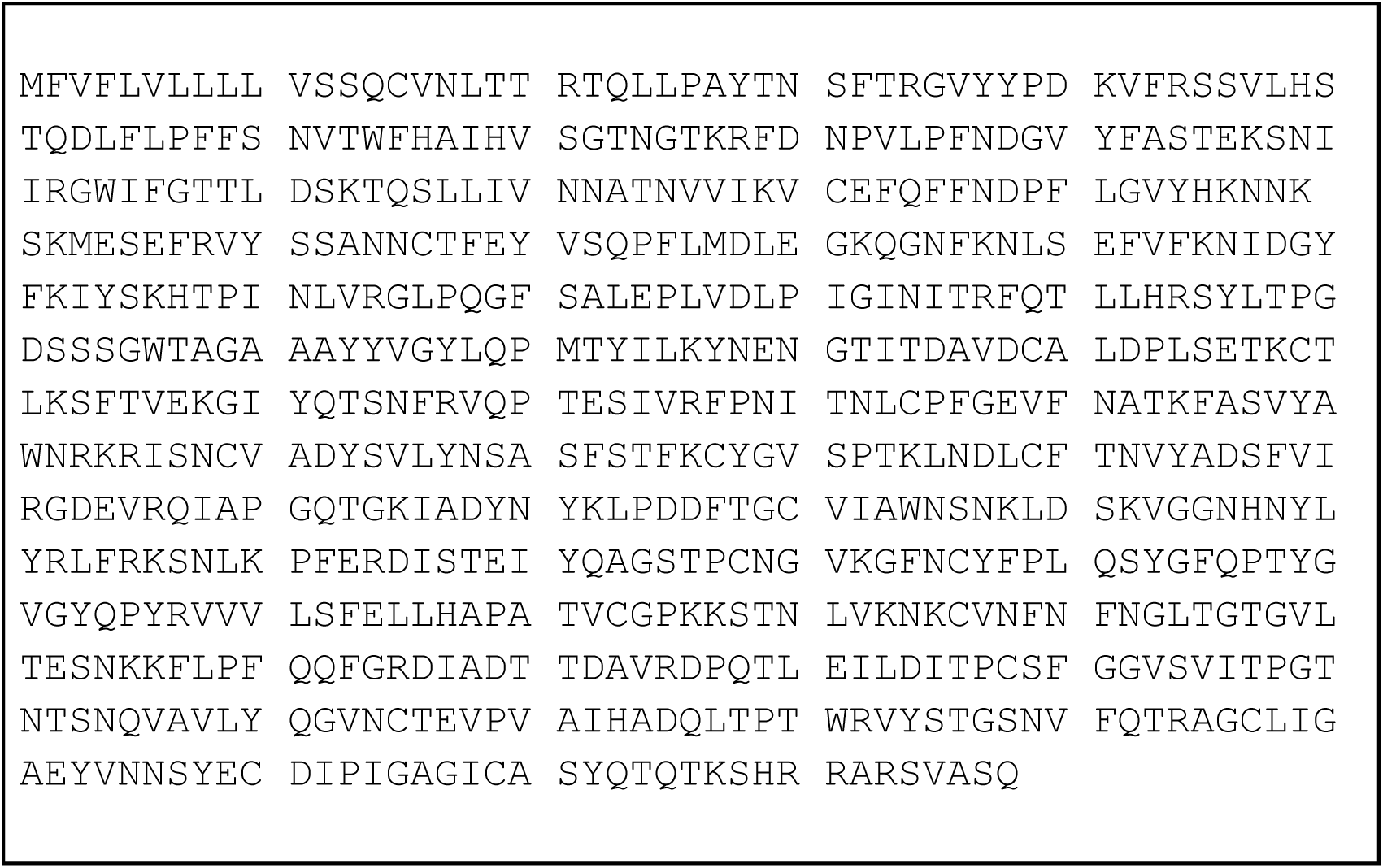
SARS-CoV-2 variant C.1.2 modified S1 spike protein, amino acids 1-687. The above sequence represents the initial construct prior to modification of the furin cleavage site and the addition of the His-tag.

**Figure 2.**
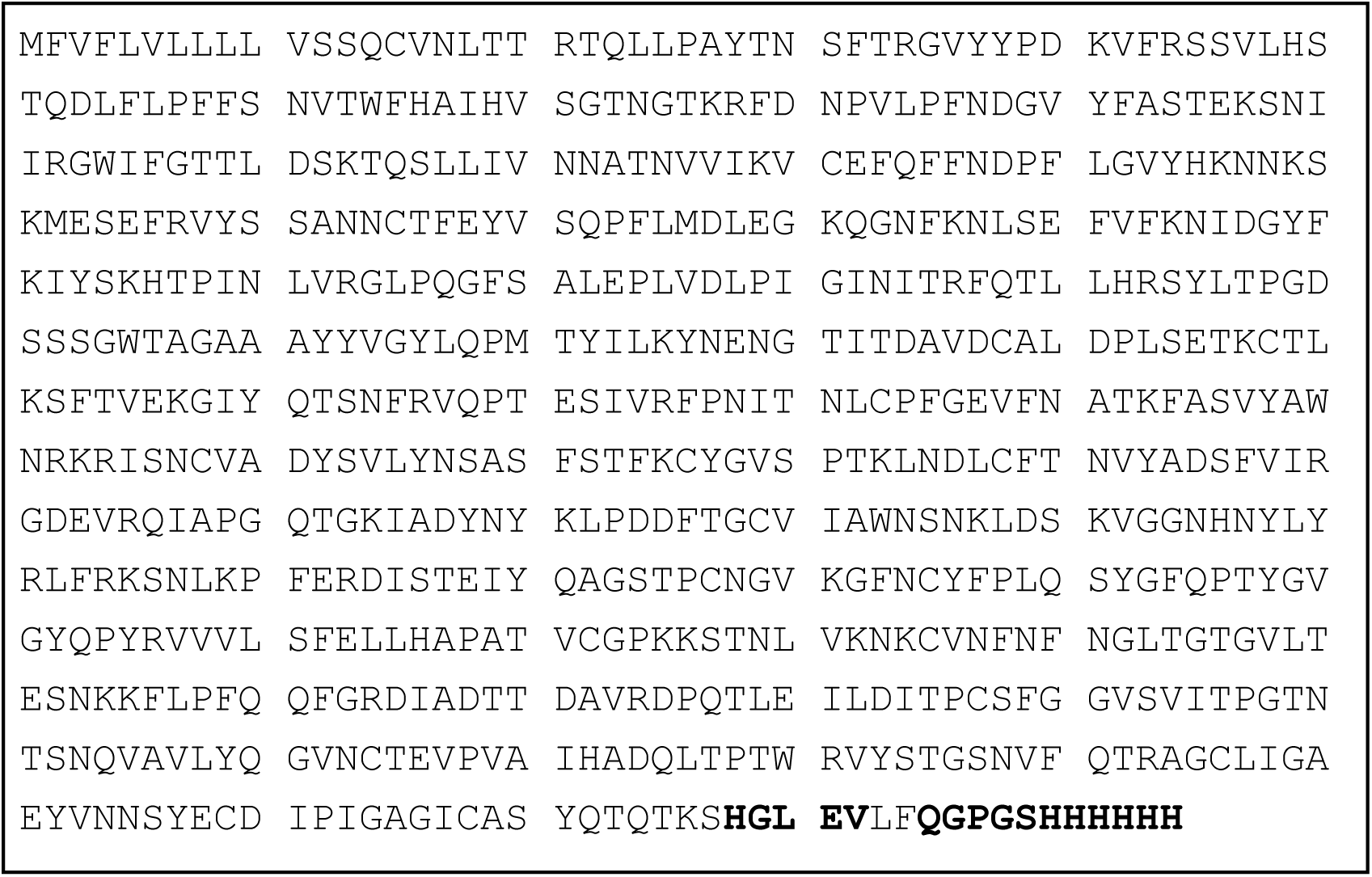
C.1.2 modified S1 Spike protein Construct (695 amino acids). Final vaccine construct is 695 amino acids of the modified SARS-CoV-2 C.1.2 variant S1 spike protein with a modification of the furin cleavage site and additional amino acid His-tag. His-tag is depicted below in black bolding at the C terminus of the construct. The furin cleavage site changes are also depicted in bold.

#### 2. Cloning of GOI from transfer vector into recombinant baculovirus

Bac-to-Bac system (ThermoFisher Invitrogen) was used for Cloning of GOI from transfer vector into recombinant baculovirus. Recombinant bacmids were produced by site-specific homologous recombination following transformation of bacmid transfer pFastBac1-GOI plasmids containing GOI into E. coli DH10Bac competent cells, which contained the AcMNPV baculovirus genome (Invitrogen). The recombinant bacmid DNA was transfected into the Sf9 insect cells seeded in 6-well plates at 0.5 × 10^6^ cells/ml using Fugene 6 reagent (Invitrogen protocol). At 72 hours post-transfection, cells were harvested for recovery in the culture medium of recombinant baculoviruses containing GOI.

#### 3. Cell culture and baculovirus infections

*Spodoptera frugiperda* Sf9 insect cells (ATCC CRL-1711) were maintained as suspension cultures in Sf900-II insect serum free medium (ThermoFisher) at 28 ◦C. Plaque isolates expressing GOI were amplified by infecting Sf9 cells seeded in shaker flasks at 2 × 10^6^ cells/ml at a multiplicity of infection (MOI)=0.05. At 72 h post-infection, culture supernatants containing the recombinant baculoviruses were harvested, clarified by centrifugation, and stored at 4 ◦C. Titers of recombinant baculovirus stocks were determined by agarose plaque assay in Sf9 cells.

#### 4. GOI Protein expression

For protein expression, Sf9 cells were infected in 200-1000 ml volume for 72 h at a cell density of 2 × 10^6^ cells/ml with recombinant baculoviruses at a MOI=3. Expression of GOI protein (Figure 2) was determined by SDS–PAGE using 4–12% gradient polyacrylamide gels (Invitrogen) and Coomassie staining and by Western blotting using antigen-specific sera.

### 2.3 Design and manufacture of adjuvanted C.1.2 S1 vaccine

The adjuvant formulations used for the above experiments were prepared using a reciprocating syringe technique which produced 5 milliliters of adjuvanted protein vaccine. The lipid formulations were composed of polyoxyethylene-2-stearyl-ether (28.10 g), cholesterol (10.8 g), Vitamin E (5.4 g), and oleic acid (120 μL), or polyoxyethylene-2-cetyl-ether (30.01 g), cholesterol (12.80 g), Vitamin E (6.00 g), and oleic acid (125 μL). SVE are the initials for the Vitamin E-containing nonphospholipid-based liposome adjuvant. On a larger scale, rotor-stator emulsion technology can be utilized inexpensively to produce large quantities of adjuvanted vaccine.

Protein was solubilized in sterile water for injection at a concentration of 1.25 mg/mL. The lipid to diluent ratio on mixing was 1:4 on a volume basis. The final concentration of protein in the adjuvanted vaccine was approximately 7 to 10 μg of protein in 268 to 356 μL of adjuvant.

Particle sizing of the exemplary vaccines was obtained via Beckman Coulter Laser Sizer LS 13 320 XR for initial samples (T1) and also samples stored at 4-8°C for one year. Stability is determined if the size of the particles does not exceed a 10% increase in size.

### 2.4 Sizing of vaccine material

Sizing of the adjuvanted vaccine product was done on the Beckman Coulter LS 13 320 XR laser sizing device. Sizing was completed at the time of manufacture and one year later. 200 μL of sample was diluted into 10.0 mL dH_2_O, loaded onto the device until the target quantity reached 6-8% and the size was measured. Data is reported as an average of two runs in nm, and the percent change reported shows product stability with a change in size of <10% increase.

### 2.5 Ethical statement for laboratory animal use and care

All animal studies were conducted at BIOQUAL in compliance with local, state, and federal regulations and were approved by BIOQUAL Institutional Animal Care and Use Committees (IACUC) protocol number 22-015. BIOQUAL, Inc. is accredited by the Association for Assessment and Accreditation of Laboratory Animal Care and in full compliance with the Animal Welfare Act and Public Health Service Policy on Humane Care and Use of Laboratory Animals.

### 2.6 Animal study design

Syrian hamsters (*Mesocricetus auratus)* were purchased from Envigo by BIOQUAL at 6 weeks of age. Upon the first immunization, hamsters were 7 weeks of age. The Syrian Golden Hamsters received a prime immunization with an average of 9.6 µg of antigen, and a booster dose with an average of 7.2 µg of antigen. Five animals were immunized subcutaneously on days 1 and 28. Serum was collected from five immunized and five non-immunized animals, and data from IgG blots for days 0, 27, 49, and 122 bleeds are shown below. A nitrocellulose dot blot technique for proteins was developed and used in this study.^9^ The proteins were the spike proteins of 2019 Wuhan-Hu-1, Delta, and Omicron BA.1, Omicron recombinant XBB.1.5, and the modified C.1.2 S1 spike protein variants of SARS-CoV-2. These proteins were utilized to detect IgG antibodies in the five animals immunized with the vaccine. The quantitative IgG response to the Wuhan-Hu-1 Receptor Binding Domain protein (Krishgen) was determined. Additionally, serum IgA and IgM antibodies were detected using the full-length Wuhan-Hu-1 spike protein, and secondary IgA and IgM antibodies (Brookwood Biomedical).

**Figure 3.**
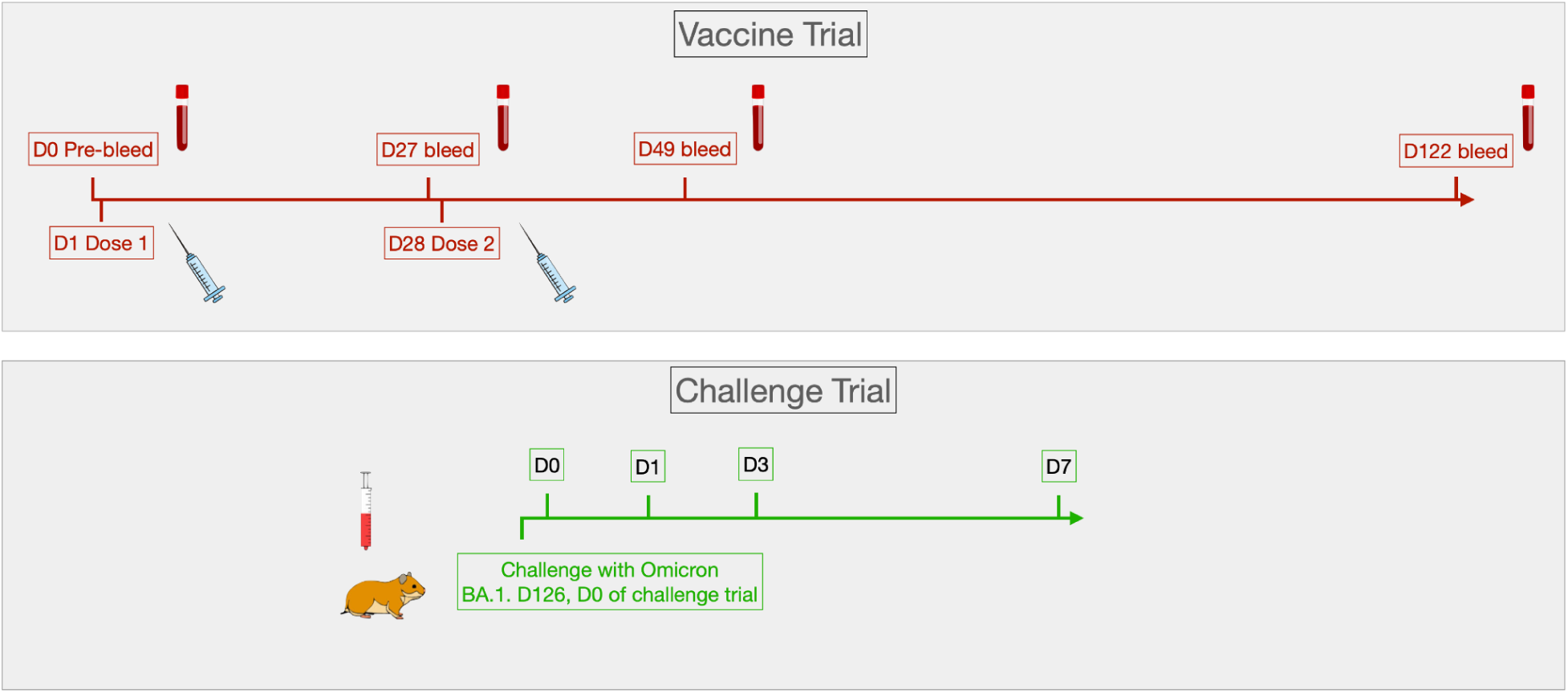
Vaccine and Challenge Trial Timeline

**Table 3.** Provides the mean amount of adjuvanted vaccine and adjuvant administered per group for Lot #021522 (n=5).

| Lot 021522 | Mean Amount of Adjuvanted Vaccine Administered per Animal (ug) | Standard Deviation | Mean Amount of Antigen Administered per Animal (ug) | Standard Deviation |
| --- | --- | --- | --- | --- |
| SVE52.C.1.2.MPS1 | 356 | 45.0555213 | 9.612 | 1.216499075 |

**Table 4.** Provides the mean amount of adjuvanted vaccine and adjuvant administered per group for Lot #031522 (n=5).

| Lot031522 | Mean Amount of Adjuvanted Vaccine Administered per Animal (ug) | Standard Deviation | Mean Amount of Antigen Administered per Animal (ug) | Standard Deviation |
| --- | --- | --- | --- | --- |
| SVE52.C.1.2.MPS1 | 268 | 49.69909456 | 7.236 | 1.341875553 |

### 2.7 Testing of sera

#### Dot blot method to detect IgG antibodies directed against the modified C.1.2 S1 protein, and full-length spike proteins of SARS-CoV-2 Wuhan-Hu-1 and variants Delta, Omicron BA.1, and Omicron recombinant XBB.1.5

Full-length spike protein antigens were purchased from Sino Biological, and the modified C.1.2 S1 protein was purchased from Medigen and stored at –20°C until utilized. Nitrocellulose sheets from BioRad were cut into 0.5×0.5 cm squares and placed in individual wells of a 12-well dish. Squares were dotted with 1 μL of 0.270 mg/mL modified C.1.2 S1 protein, or 0.25 mg/mL of full-length spike protein antigen solution and then allowed to dry for 30 minutes at room temperature (RT). Squares were then blocked in 1 mL Blocker buffer (1% milk protein (Thermo Scientific)) in phosphate buffered saline, pH=7.4 (Gibco) for 1 hour at RT on a plate rocker. Animal serum was diluted at 1:500 by adding 2 μL serum into 998 μL Blocker. This was then added to each well and allowed to incubate for 2 hours at RT on a plate rocker. For dilution experiments, 1:2 dilutions were performed in 15 mL sterile conical tubes prior to addition to plates. The dot blots were washed twice with 2 mL phosphate buffered saline (pH=7.4) for 4 minutes each. Secondary antibody conjugated to alkaline phosphatase was added in Blocker at a dilution of 1:100 to each well and incubated for 1 hour at RT on a plate rocker. Developing solution was made by adding 10 mg Naphthol AS_MX phosphate disodium salt (Sigma Aldrich) and 22 mg Fast Red TR salt (Sigma Aldrich) into 10 mL Tris-HCL pH=8.0 (Sigma Aldrich). Blots were then washed twice with 4 mL PBS for 4 minutes each and then again with 4 mL Tris-HCL pH=8.0. Nitrocellulose was developed by adding 1 mL developing solution. A positive result occurs when a red dot appears on the nitrocellulose. Dot blots were then washed with 2 mL of PBS for 2 minutes and allowed to dry overnight at RT. Photographs of the nitrocellulose were taken the next day. All animals were immunized subcutaneously.

#### Quantitative determination of hamster anti-RBD-SARS-CoV-2 IgG antibody levels

Antibody levels were determined using the Krishgen Bioscience GENLISA Hamster Anti-SARS-CoV-2 (Covid-19) IgG Antibody to Spike RBD protein Quantitative Titration ELISA kit. Kit reagents were brought to room temperature prior to use. Antibody standard was reconstituted at a concentration of 1000 ng/mL in the provided standard diluent. Serum samples of individual animals were diluted from 1:1000 to 1:64,000 by performing 1:2 serial dilutions in the provided sample diluent. Standard was diluted as a seven point concentration curve with the following concentrations; 720 ng/mL, 360 ng/mL, 180 ng/mL, 90 ng/mL, 60 ng/mL, 30 ng/mL, and 15 ng/mL. 100 uL of sample and standard were added to individual wells that came pre-coated with antigen. Plate was sealed and incubated for 1 hour at room temperature (RT). Plate was then washed 4 times with 100 uL of 1× Wash Buffer. 100 uL of goat anti-hamster IgG HRP conjugated secondary antibody was added and the plate was sealed and incubated at RT for 1 hour. Plate washed 4 times with 100 uL of 1× Wash Buffer. 100 uL of TMB substrate was added to each well and the plate was incubated in the dark at RT for 15 minutes. Reaction stopped by the addition of 100 uL Stop Solution. The plate was read at 450 nm on the Byonoy Absorbance 96 plate reader. Second order polynomial nonlinear regression used to generate the standard curve, and concentration of serum samples were determined by interpolating from the standard curve.

#### Dot blot method to detect IgA and IgM antibodies directed against the spike protein of SARS-CoV-2 Hu-1

Spike protein antigens were purchased from Sino Biological and stored at –20°C until utilized. Nitrocellulose sheets from BioRad were cut into 0.5×0.5 cm squares and placed in individual wells of a 12-well dish. Squares were dotted with 1 μL of 0.25 mg/mL antigen solution and then allowed to dry for 30 minutes at room temperature (RT). Squares were then blocked in 1 mL Blocker buffer (1% casein protein (Thermo Scientific) in phosphate buffered saline for 1 hour at RT on plate rocker. Animal serum was diluted at 1:500 by adding 2 μL serum into 998 μL Blocker. This was then added to each well and allowed to incubate for 2 hours at RT on a plate rocker. For dilution experiments, 1:2 dilutions were performed in 15 mL sterile conical tubes prior to addition to plates. The dot blots were washed twice with 2 mL phosphate buffered saline (pH=7.4, Gibco) for 4 minutes each. Secondary IgM or IgA antibody conjugated to alkaline phosphatase (Brookwood Biomedical) was added in Blocker at a dilution of 1:100 to each well and incubated for 1 hour at RT on a plate rocker. Developing solution was made by adding 10 mg Naphthol AS_MX phosphate disodium salt (Sigma Aldrich) and 22 mg Fast Red TR salt (Sigma Aldrich) into 10 mL Tris-HCL pH=8.0 (Sigma Aldrich). Blots were then washed twice with 4 mL PBS for 4 minutes each and then again with 4 mL Tris-HCL pH=8.0. Nitrocellulose was developed by adding 1 mL developing solution. A positive result occurs when a red dot appears on the nitrocellulose. Dot blots were then washed with 2 mL of PBS for 2 minutes and allowed to dry overnight at RT. Photographs of the nitrocellulose were taken the next day.

### 2.8 qRT-PCR assay

BIOQUAL, Inc. (Gaithersburg, MD) performed the animal study and collected weights, clinical observations, sera, and swabs for PCR assays. The qRT-PCR assay was performed at UC Davis under a BIOQUAL subcontract.

#### Oral swabs collection

The animal was restrained properly for sample collection. A sterile flocked FLOQ swabs (COPAN Cat. #: 501CS01) was removed from the packaging. Each hamster was swabbed on the inside of the mouth and both cheeks with one swab for at least 30 seconds. The swab was then placed into a cryovial with 1 mL PBS, the shaft cut off to fit into the vial. The oral swab sample was then immediately snap-frozen on dry ice and stored at –80°C until viral load analysis.

#### Quantitative RT-PCR assay for SARS-CoV-2 RNA in oral swabs

Viral load analysis of oral swab samples was performed at UC Davis. Viral RNA was extracted from oral swab samples by homogenization in TRIzol LS and phase separation using BCP. RNA was extracted from the aqueous phase and purified using RNeasy minikits (Qiagen). Following DNase treatment (ezDNAse; Invitrogen), cDNA was generated using super-script IV reverse transcriptase (Thermo Fisher) with RNAseOUT (Invitrogen). cDNA from samples was mixed with QuantiTect probe PCR master mix and gene-specific primers to run on a QuantStudio 6 Flex real-time cycler (Applied Biosystems). To generate a control for the amplification reaction, oligos of the amplicon template were used and the copy number calculated based on the mass and RNA molar mass (340.5 g/mole). Standard curve was prepared by 10-fold serial dilution to give a range of 1 to 10^7^ copies per reaction and tested by qPCR. The number of RNA copies per mL for samples was calculated by extrapolation from the standard curve and volume corrected to give RNA copies per mL starting sample volume. The qRT-PCR assay for SARS-CoV-2 genomic RNA (gRNA) utilized primers and a probe specifically designed to amplify and bind to a conserved region of Nucleocapsid (N) gene of SARS-CoV-2. The qRT-PCR assay for SARS-CoV-2 subgenomic RNA (sgRNA) utilized primers and a probe specifically designed to amplify and bind to a region of the Envelope (E) gene messenger RNA from SARS-CoV-2, which is not packaged into the virion.

#### Primers/probe sequences for gRNA amplification

2019-nCoV Forward Primer: 5’-GAC CCC AAA ATC AGC GAA AT-3’

2019-nCoV Reverse Primer: 5’-TCT GGT TAC TGC CAG TTG AAT CTG-3’

2019-nCoV P: 5’-FAM-ACC CCG CAT /ZEN/ TAC GTT TGG TGG ACC-BHQ1-3’

#### Primers/probe sequences for sgRNA amplification

Sg-E Forward Primer: 5’-CGA TCT CTT GTA GAT CTG TTC TC –3’

E Sarbeco R2 Reverse Primer: 5’-ATA TTG CAG CAG TAC GCA CAC A-3’

E Sarbeco P1: 5’-/FAM-ACA CTA GCC ATC CTT ACT GCG CTT CG-BBQ –3’

### 2.9 Clinical observations

Hamsters were observed daily for overall health conditions during the study. Body weights were recorded at the time of vaccination and then daily after challenge. The baseline body weights of all the animals were measured before virus infection. After challenge, animals were monitored for 7 consecutive days for clinical signs such as weight loss, lethargy, ruffled fur, hunched back, labored breathing, and ocular discharge.

### 2.10 Statistical analysis

No statistical plan was outlined in the original animal study for the quantitative antibody determination. A post-hoc paired t-test analysis of the quantitative assays was conducted at days 0, 27, 49, and 122. Data were plotted using GraphPad Prism (GraphPad software, version 8.0.0; https://www.graphpad.com).

## 3. Results

### 3.1 Sizing and stability

**Table 5.**
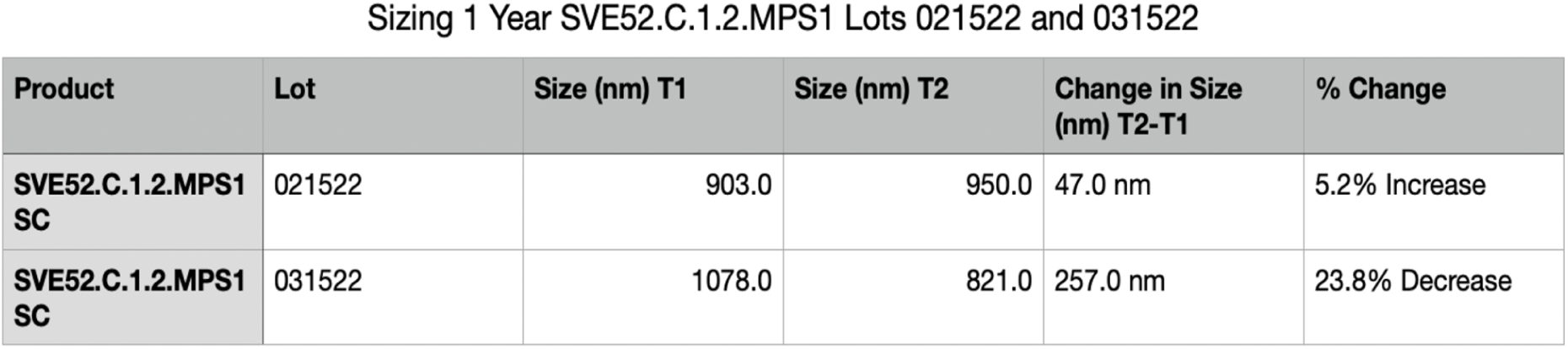
Particle sizing results for vaccines SVE52.C.1.2.MPS1 from two lots of vaccine production (Lots #021522 and #031522). Lots #021522 and #031522 sizing was done initially and one year later. 200 μL of sample was diluted into 10.0 mL dH2O, loaded onto the device until the target quantity reached 6-8% and the size was measured. Data is reported as an average of two runs in nm, and the percent change reported shows product stability with a change in size of <10% increase.

### 3.2 Serologic testing

#### Serum IgG antibody responses to Wuhan-Hu-1 full-length spike protein

The immunogenicity of SVE52.C.1.2.MPS1 administered to hamsters twice by subcutaneous inoculation with C.1.2 modified S1 spike protein was measured by dot blot (Figures 4-6). We have demonstrated that, after one dose of vaccine, five out of five animals were positive on day 27 to the following antigens: modified C.1.2 S1 protein, full-length Wuhan-Hu-1, and Delta. A booster dose was given on day 28. At day 49 and day 122, IgG antibody responses to all antigens were noted. Sera IgG antibodies were also tested against the full-length spike proteins of Omicron BA.1 and the recombinant Omicron XBB.1.5, with three animals showing positive results for BA.1, and four animals showing positive results for XBB.1.5 after the first dose. Bleeds at days 49 and 122 showed a positive response for all five animals for both antigens. The blots for Omicron BA.1 and recombinant Omicron XBB.1.5 are not shown.

**Figure 4.**
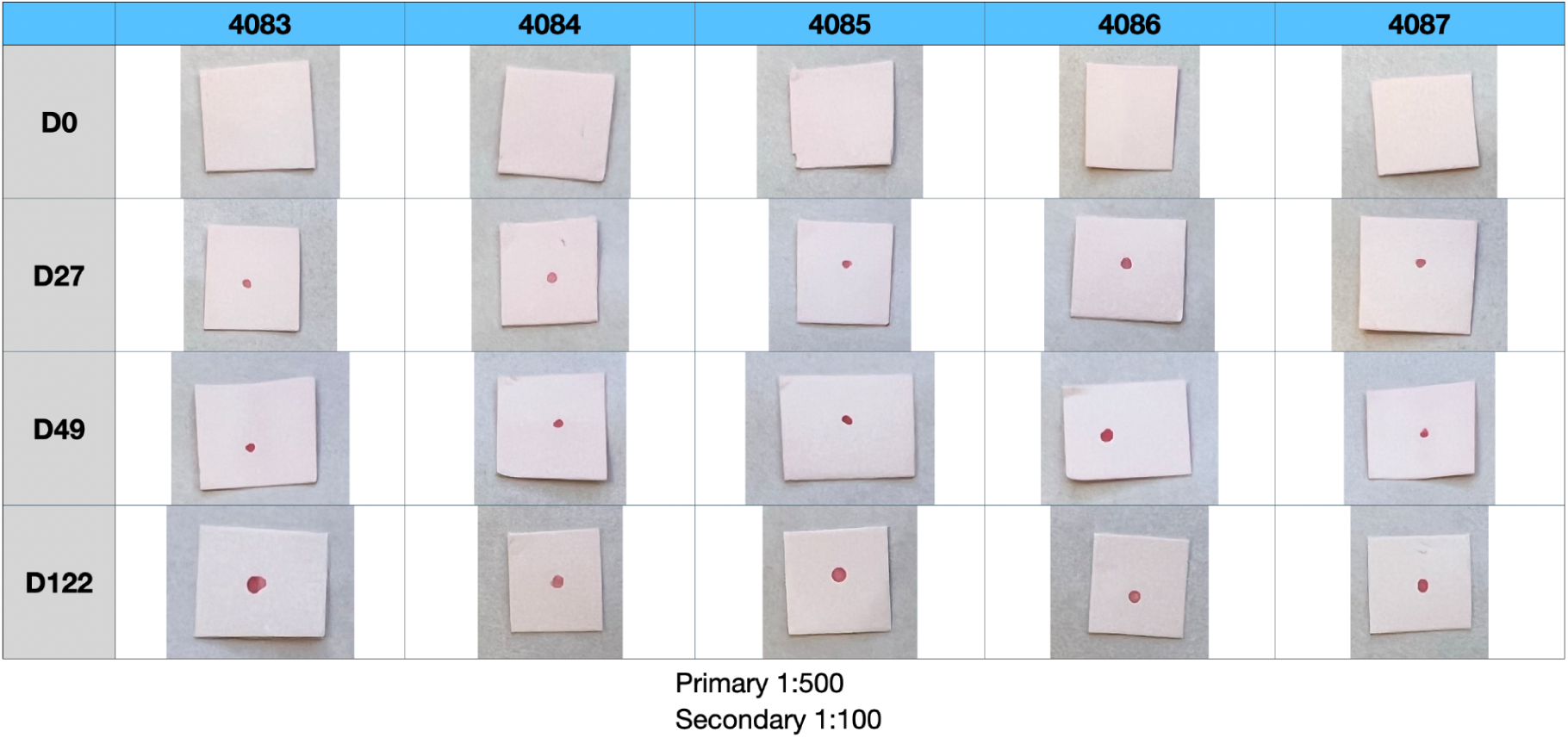
Depicts blot images for IgG antibodies directed against the modified S1 spike protein of the C.1.2 isolate. The animals were subcutaneously immunized with a SVE52 nonphospholipid liposome (containing vitamin E) and a modified S1 sequence of the C.1.2 variant of SARS-CoV-2 (Figure 2). Animals were bled at days 0, 27, 49, and 122. Animals were immunized on days 1 and 28. The dilution of the sera was 1:500. High antibody responses by blot were noted in all five animals at days 27, 49, and 122.

**Figure 5.**
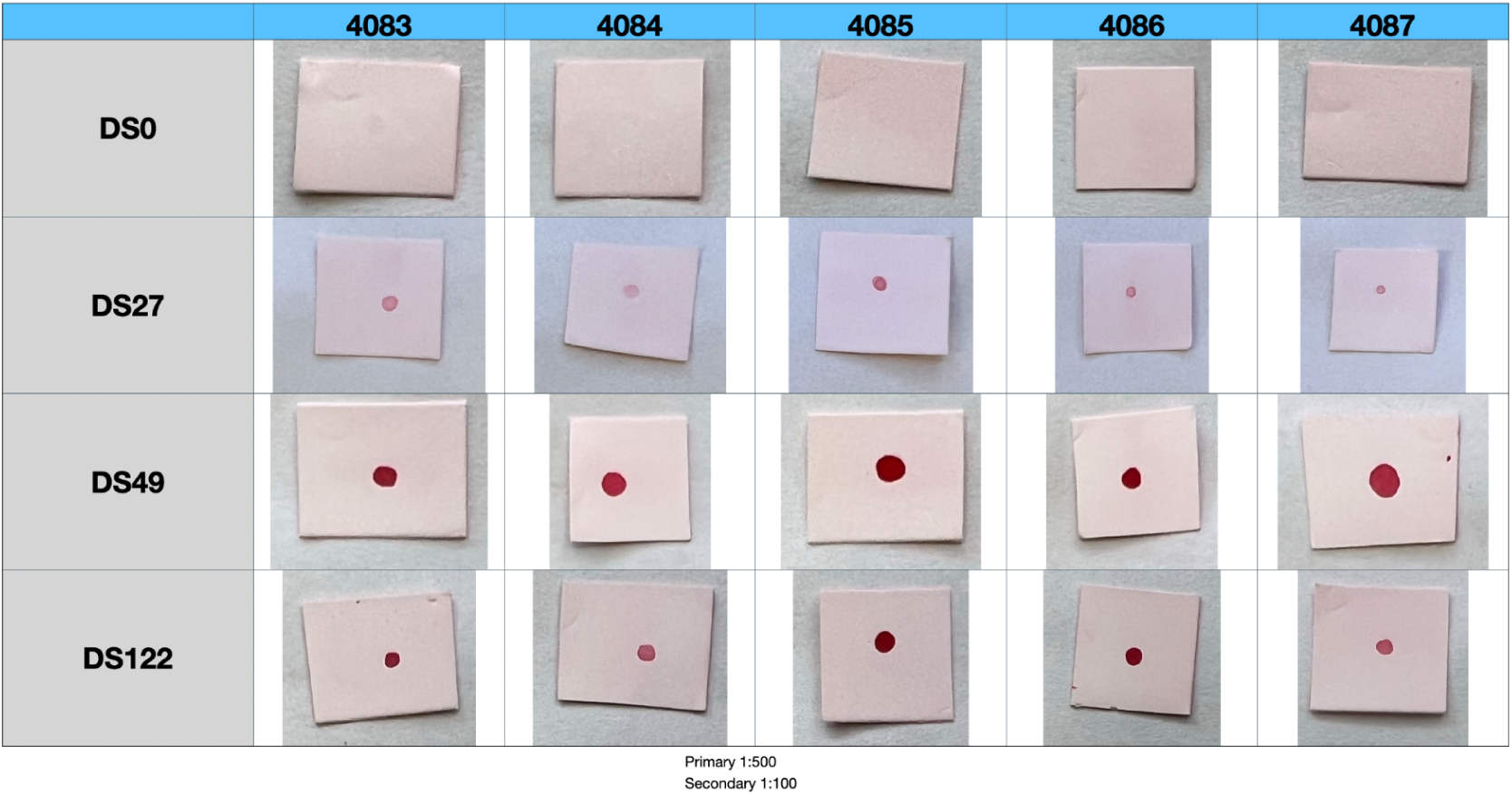
Depicts blot images for IgG antibodies directed against the full-length spike protein of the 2019 Wuhan-Hu-1 SARS-CoV-2 isolate. The animals were subcutaneously immunized with a SVE52 nonphospholipid liposome (containing vitamin E) and a modified S1 sequence of the C.1.2 variant of SARS-CoV-2 (Figure 2). Animals were bled at days 0, 27, 49, and 122. Animals were immunized on days 1 and 28. The dilution of the sera was 1:500. High antibody responses by blot were noted in all five animals at days 27, 49, and 122.

**Figure 6.**
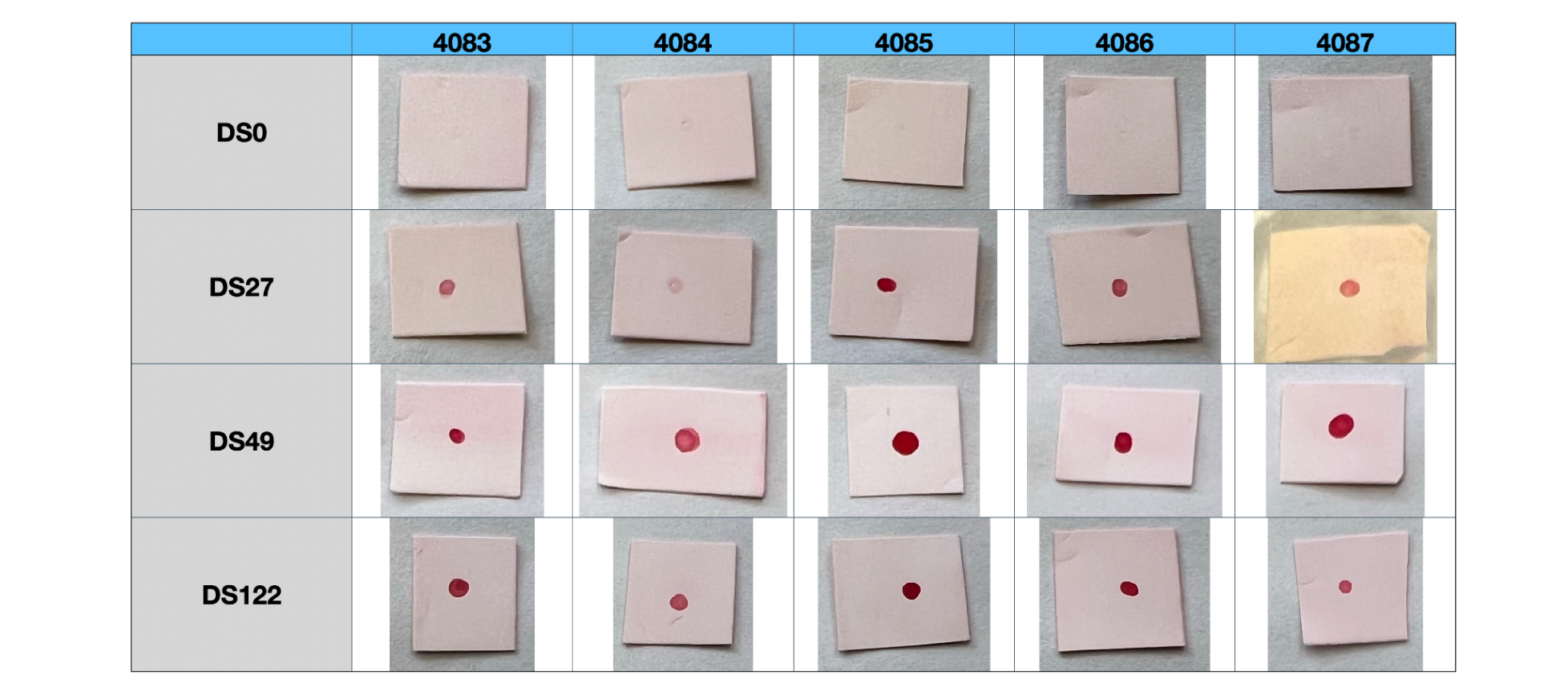
Depicts blot images for IgG antibodies directed against the full-length spike protein of the SARS-CoV-2 Delta B.1.617.2 variant. The animals were subcutaneously immunized with a SVE52 nonphospholipid liposome (containing vitamin E) and a modified S1 sequence of the C.1.2 variant of SARS-CoV-2 (Figure 2). Animals were bled at days 0, 27, 49, and 122. Animals were immunized on days 1 and 28. The dilution of the sera was 1:500. High antibody responses by blot were noted in all five animals at days 27, 49, and 122.

#### Quantitative analysis of IgG antibodies to the Receptor Binding Domain

Quantitative IgG antibody analysis determined with the Krishgen ELISA kit shows a positive response in three out of five animals at day 27. Quantitative analysis at day 49 shows an anamnestic response in five out of five animals. At day 122, 94 days after the boost, five out of five animals still had measurable levels of IgG antibody to the Wuhan-Hu-1 Receptor Binding Domain.

**Figure 7.**
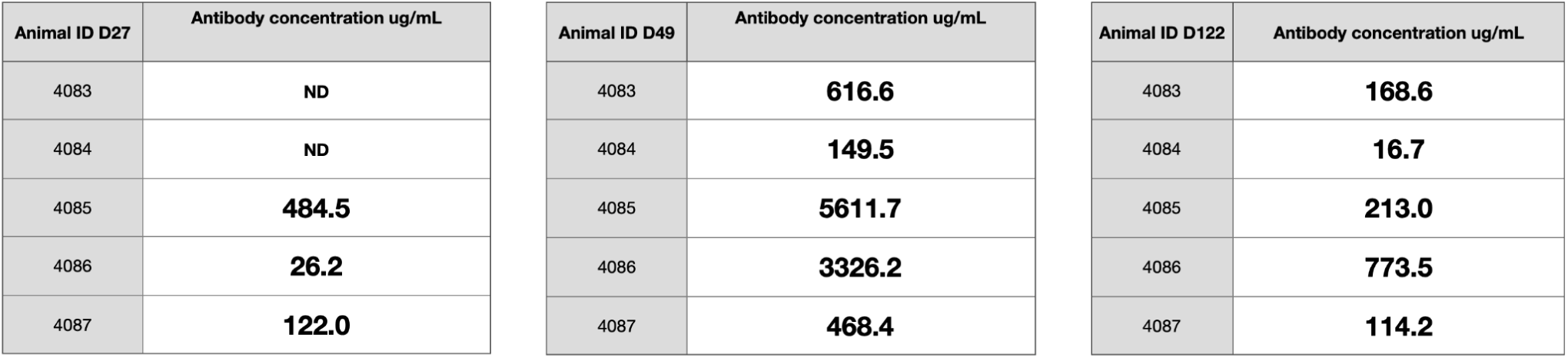
Describes the quantitative IgG antibody response in hamsters to Wuhan-Hu-1 RBD as measured by the Krishgen BioSystems GENLISA Hamster Anti-SARS-CoV-2 IgG Antibody Quantitative Titration ELISA kit at days 27, 49, and 122. Quantifiable IgG antibody levels were measured at day 27 in three out of five animals after one subcutaneous dose of vaccine. An anamnestic response was observed on day 49 after the second dose was administered on day 28. All five out of five animals had a quantifiable antibody response that persisted to day 122, with a decline in antibody response from the day 49 measurement. Day 27 one-tail p value=0.12. Day 49 one-tail p value=0.06. Day 122 one-tail p value=0.06.

#### Serum IgM and IgA antibody responses to Wuhan-Hu-1 full-length spike protein

Apart from IgG, IgM and IgA are also critical components of the immune system. Reagents became available to us which allowed us to look at IgM and IgA responses to full-length spike protein from the Wuhan-Hu-1 isolate. We found positive serum IgM and IgA responses in five out of five animals at days 27, 49, and 122.

**Table 6.**
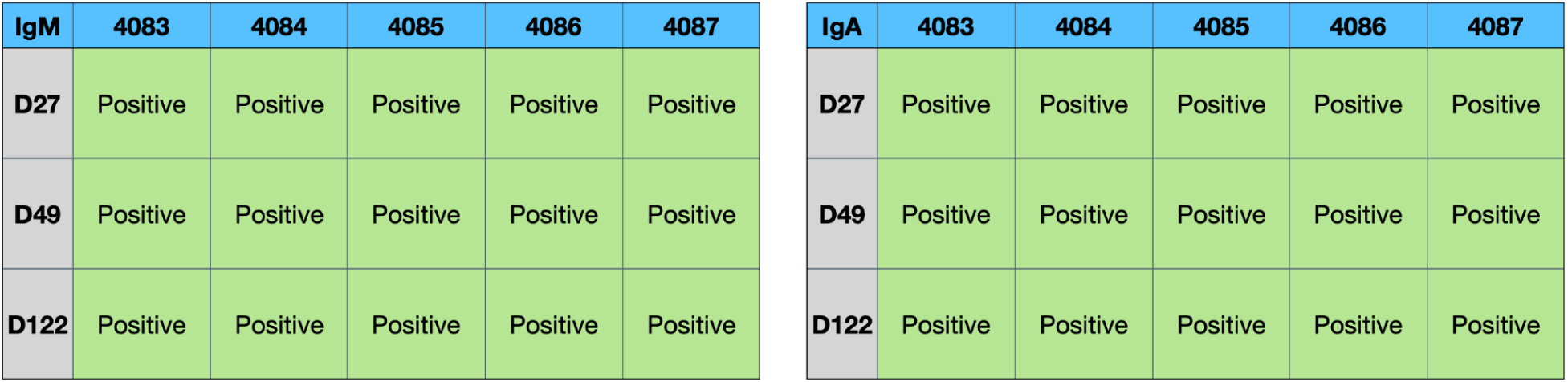
Qualitative dot blot measuring IgM and IgA antibodies to the Wuhan-Hu-1 full-length spike protein shows a positive response on day 27 after one dose of subcutaneous vaccine in five out of five animals. The IgM and IgA antibody responses were positive on day 49 after the second dose, with persistent antibody responses measured out to day 122.

| IgM | 4083 | 4084 | 4085 | 4086 | 4087 |
| --- | --- | --- | --- | --- | --- |
| **D27** | Positive | Positive | Positive | Positive | Positive |
| **D49** | Positive | Positive | Positive | Positive | Positive |
| **D122** | Positive | Positive | Positive | Positive | Positive |

| IgA | 4083 | 4084 | 4085 | 4086 | 4087 |
| --- | --- | --- | --- | --- | --- |
| **D27** | Positive | Positive | Positive | Positive | Positive |
| **D49** | Positive | Positive | Positive | Positive | Positive |
| **D122** | Positive | Positive | Positive | Positive | Positive |

### 3.3 Live challenge study in Syrian hamsters subcutaneously immunized with modified C.1.2 S1 protein vaccine

#### Clinical observations

After challenge with the Omicron BA.1 variant, clinical observations were made and body weights were measured every day for seven days. Five out of five animals were successfully infected with the challenge strain. All animals lost weight through day 6. By day 7, animals in the treated group began to gain weight.

**Figure 8.**
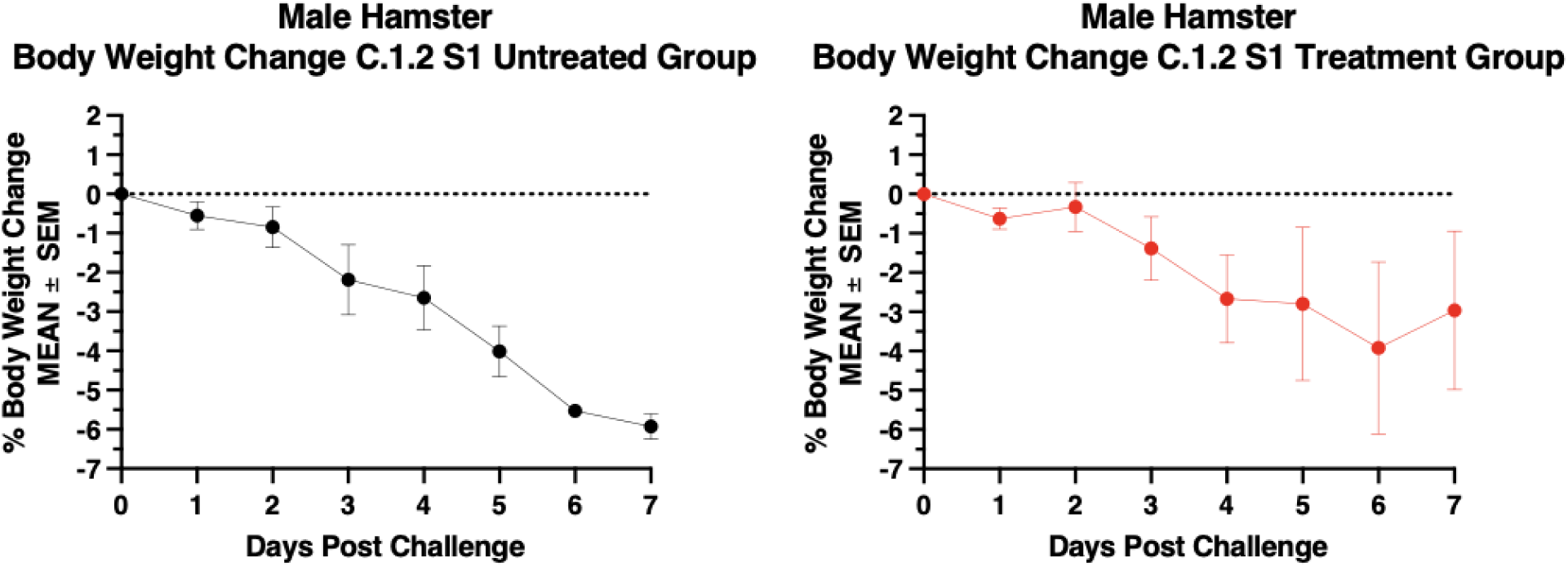
Depicts the percent of body weight change upon a live viral challenge with the Omicron BA.1 isolate. Animals were challenged on day 0 of the challenge trial (n=5 per group). Body weights and clinical observations were taken every day for 7 days post-challenge. Body weights in the control group show a greater decrease in percent body weight compared to the treated group. The treated group shows an increase in body weight between days 6 and 7, indicating a protective effect upon vaccination. Error bars represent ±SEM.

#### Quantitative RT-PCR assay for SARS-CoV-2 RNA in oral swabs

The quantitative RT-PCR assay for SARS-CoV-2 RNA in oral swabs was performed at the UC Davis National Primate Center. Animals in the untreated group at day 1 and day 2 had similar amounts of detected virus. By day 3, a decrease in viral load was noted in the vaccine group. By day 7, five out of five animals in the vaccine group had undetectable levels of virus, while only two out of five animals in the untreated group had undetectable levels of virus. Because of the small number of animals in each group, statistical analysis on this data set is difficult.

**Figure 9.**
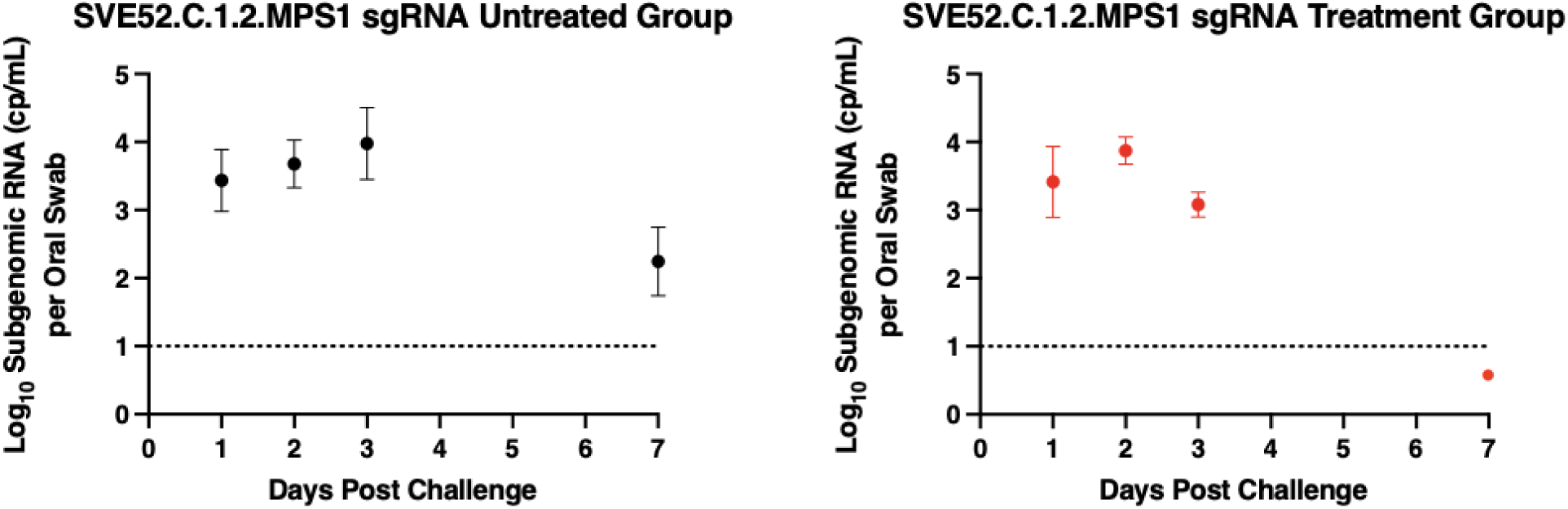
Depicts the viral subgenomic RNA qRT-PCR results of the treated group and the untreated group (n=5 per group) following a challenge with Omicron BA.1 on day 0. Oral swabs were taken at days 1, 2, 3, and 7 post infection. Data points below the limit of detection indicating no virus detected are shown as Log_10_(0.5). Immunized animals begin to show a decrease in viral titer by day 3, with one animal in the treatment group being below the limit of detection and five animals in the untreated group having detectable virus. By day 7, no animals in the treatment group and three animals in the untreated group had no detectable virus. Data points are represented as mean of viral load in detectable animals. Error bars represent ±SD.

## 4. Discussion

Highly effective vaccines against SARS-CoV-2 have been developed, tested and approved in the U.S. with U.S. government funding. Many of these vaccines require a complicated cold chain (Moderna, Pfizer) or use live adenoviral vectors (Johnson & Johnson). All of these require IM injections and have not produced long-lasting immunity.

At the start of the COVID-19 pandemic, we designed a modified S1 protein based on the C.1.2 variant and contracted Medigen to produce the baculovirus-expressed modified SARS-CoV-2 spike protein. We have spent two years evaluating the hypothesis that we could make inexpensive adjuvants for SARS-CoV-2 spike proteins that could be administered in a one or two-dose format subcutaneously. These adjuvanted vaccines have demonstrated stability for at least a year and could be stored at 2-8°C. In these vaccines, we deleted sequences previously identified by Lyons-Weiler as having cross-reactivity with known human peptide protein sequences. The construct we describe in this small, preliminary animal study is a modified S1 sequence of the C.1.2 variant of SARS-CoV-2 encapsulated in a nonpospholipid liposome vaccine.

We have demonstrated that a modified S1 vaccine injected subcutaneously generates IgG, IgM, and IgA antibody responses after a single dose, with responses increasing after a second dose. Moreover, this vaccine provides prolonged antibody responses to Wuhan-Hu-1, Delta, Omicron BA.1, and the recombinant Omicron XBB.1.5 through at least day 122. In addition, our vaccine demonstrated a notable reduction in disease severity when measured by weight loss and viral titer. Vaccinated hamsters had a decreased rate of weight loss compared to the untreated group over the course of seven days, with a weight gain noted between days 6 and 7. Vaccinated hamsters also had a decrease in viral load compared to the untreated group, with all vaccinated hamsters showing no detectable virus by day 7.

The drawbacks to our initial study with this adjuvanted modified S1 spike protein vaccine are that there were only 5 animals in each group. Another issue with our vaccine is that we have not conducted a dose response experiment.

In summary, the vaccine’s protective impact persisted 98 days after the second immunization, providing long-term immunogenicity to notable variants of concern, and suggesting the potential for a long-term protective effect. These findings underscore the potential of this adjuvant technology for use in the development of a viable, long-term, subcutaneously administered vaccine.

## Author Contributions

Dr. David Craig Wright conceived the project and conducted vaccine and adjuvant design. Vaccine manufacturing was completed by Dr. David Craig Wright and Michael Bowe. Michael Bowe performed all assays done in house, including dot blot analysis, ELISA assays, and vaccine sizing. Dr. Backstedt of BIOQUAL was the study lead for the animal trial. Casey Gardner of BIOQUAL acted as the Animal Project Manager and previously during the study was the technician performing immunization and other in-life procedures. Micah Short-Freeman of BIOQUAL was active in the viral challenge preparations. Dr. Swagata Kar was also involved with the study. Dr. Pushko supervised antigen production. Dr. Wright, Emily Wright, and Michael Bowe drafted and revised the manuscript. All authors reviewed the data.

## Abbreviations

ACE2: Angiotensin-converting enzyme 2
BSL: Biosafety level
COVID-19: Coronavirus disease 2019
Dpi: Days post infection
ELISA: Enzyme-linked immunosorbent assay
G: Gram
GOI: Gene of Interest
IgG: Immunoglobulin G
MOI: Multiplicity of Infection
IgM: Immunoglobulin M
IgA: Immunoglobulin A
RBD: Receptor-binding domain
S: Spike or surface glycoprotein
SARS-CoV-2: Severe acute respiratory syndrome coronavirus 2
sc: subcutaneous
ND: Not Detected
SVE52.C.1.2.MPS1: negatively-charged nonphospholipid-based liposome containing vitamin E and the modified S1 spike protein
XBB.1.5: this SARS-CoV-2 subvariant is a sublineage of the XBB variant, a recombinant of two BA.2 sublineages

## Acknowledgements

We thank Chris Miller and Linda Fritts of the California National Primate Research Center (CNPRC) at UC Davis for technical assistance conducting oral swab PCR analysis. We also thank Sean Court of Medigen, Inc. for C.1.2 protein purification. Dr. Wright would also like to recognize Edward B. Hager, M.D. (deceased) and Donald F. H. Wallach, M.D. (deceased), with whom he worked for many years.

